# Large increases in emissions of methane and nitrous oxide from eutrophication in Lake Erie

**DOI:** 10.1101/648154

**Authors:** Julianne M. Fernandez, Amy Townsend-Small, Arthur Zastepa, Susan B. Watson, Jay A. Brandes

## Abstract

Eutrophication is linked to greenhouse gas emissions from inland waters. Phytoplankton blooms in Lake Erie, one of Earth’s largest lakes, have increased with nutrient runoff linked to climate warming, although greenhouse gas emissions from this or other large eutrophic lakes are not well characterized. We measured greenhouse gases around Lake Erie in all four seasons and found that CH_4_ and N_2_O emissions have increased 10 times or more with re-eutrophication, especially during and after phytoplankton blooms. Lake Erie is a positive source of CH_4_ throughout the entire year and around the entire lake, with the highest emissions in spring and summer near the mouth of the Maumee River. While Lake Erie is an overall N_2_O source, it is an N_2_O sink in winter throughout the lake and in some locations during large phytoplankton blooms. We estimate that Lake Erie emits ~6300 metric tons of CH_4_-C yr^−1^ (± 19%) and ~600 metric tons N_2_O-N yr^−1^ (± 37%): almost 500,000 metric tons CO_2_-eq yr^−1^ total. These results highlight the gravity of eutrophication-related increases in large lake GHG emissions: an overlooked, but potentially major feedback to global climate change.

## Introduction

Lake Erie is one of Earth’s largest eutrophic lakes, with a history of eutrophication linked to decades of urban, agricultural, and industrial impacts (1). The Great Lakes Water Quality Agreement in 1972 reduced point source nutrient pollution and phytoplankton biomass (2). However, since the mid-1990s, Lake Erie has experienced “re-eutrophication”, especially the Western Basin, attributed to a rise in rainfall and non-point source fertilizer runoff from the Maumee River watershed (2)(3). This has led to massive, sometimes toxic, phytoplankton blooms, drinking water intake closures, and exacerbation of central basin hypoxia (3)(4)(5). Climate change is implicated in water quality declines, as increased intensity and rates of precipitation increases watershed nutrient and sediment inputs (6).

Lakes, rivers, and reservoirs play a major role in the global carbon cycle and greenhouse gas emissions budgets (7–11). Inland waters are a major source of CH_4_ globally, and anthropogenic construction of reservoirs enhances this flux (12). Eutrophication leads to CH_4_ and N_2_O evasion from lakes (13)(14)(10), and eutrophication will increase in lake and coastal ecosystems as runoff and agricultural activity increase globally (6)(15). The greenhouse gases CH_4_ and N_2_O are 25 and 289 times more powerful than carbon dioxide (CO_2_) over 100 year time scales, respectively (16). Despite decades of research on nutrients and phytoplankton in the Great Lakes, there is sparse research on GHGs in Lake Erie (17)(18)(19)(20)(21). However, if increasing eutrophication in Lake Erie linked to climate change leads to increasing emissions of CH_4_ and N_2_O, this represents a positive feedback that urgently needs to be both delineated and addressed (22).

Lake Erie is the shallowest in average depth (~19 m), has the shortest water residence time, and is the lowest in latitude of the North American Great Lakes (23,24). The lake is divided lengthwise by an international border between the northern United States and the most southern portion of Canada. The lake is also roughly divided into three main basins by depth: the Western Basin, the Central Basin, and the Eastern Basin. The watershed is urbanized, industrialized, and agricultural, with 17 cities in both countries including several industrial hubs (24,25). The main water source into Lake Erie is the upper Great Lakes via the Detroit River (24). Although drilling for oil and gas in the Great Lakes is currently banned in the United States, the Canadian waters of Lake Erie have been actively drilled for natural gas by Canada since the 1930s (18). Currently, there are over 2000 conventional natural gas wells offshore in the Canadian waters of Lake Erie (http://www.ogsrlibrary.com), approximately 500 of which are actively producing. The lake is characterized by high cyanobacteria and phytoplankton biomass, including the harmful cyanobacteria *Microcystis*, in the Western basin (and to a lesser extent, the Central basin), driven by spring and summer nutrient runoff from the Maumee and Sandusky Rivers (2,6). Past research from our research group has indicated Lake Erie is a positive source of CH_4_ to the atmosphere in the Central Basin in the late summer season (18).

In order to determine whether Lake Erie is an annual source of CH_4_ and N_2_O over the entire surface, we carried out CH_4_ and N_2_O measurements throughout Lake Erie during all four seasons from May 2015 – August 2016, including two mid-summer dates (August 2015 and 2016) and one in winter (February 2016), when the lake had partial ice cover. We also made measurements of CO_2_ emissions and sources (using stable isotopes) during August 2016.

## Materials and methods

### Sample collection

Sampling was conducted aboard the Canadian Coast Guard Ship (CCGS) *Limnos* and the CCGS *Griffon*. Water was collected from the Western, Central, and Eastern basin of the lake, with sampling conducted on both sides of the international border. Cruise dates were as follows: 12^th^ - 15^th^ May 2015, 24^th^ - 27^th^ August 2015, 5^th^ - 8^th^ October 2015, 15^th^ - 19^th^ February 2016, and 29^th^ August – 1^st^ September 2016 (Fig 1, Tables S1-S6). A YSI EXO 2 Multi-parameter Sonde, equipped with an EXO Conductivity and Temperature Smart Sensor, EXO Optical Dissolved Oxygen Smart Sensor, and a guarded EXO pH Smart Sensor, was deployed at each station for water column profile characteristics.

**Fig. 1.**
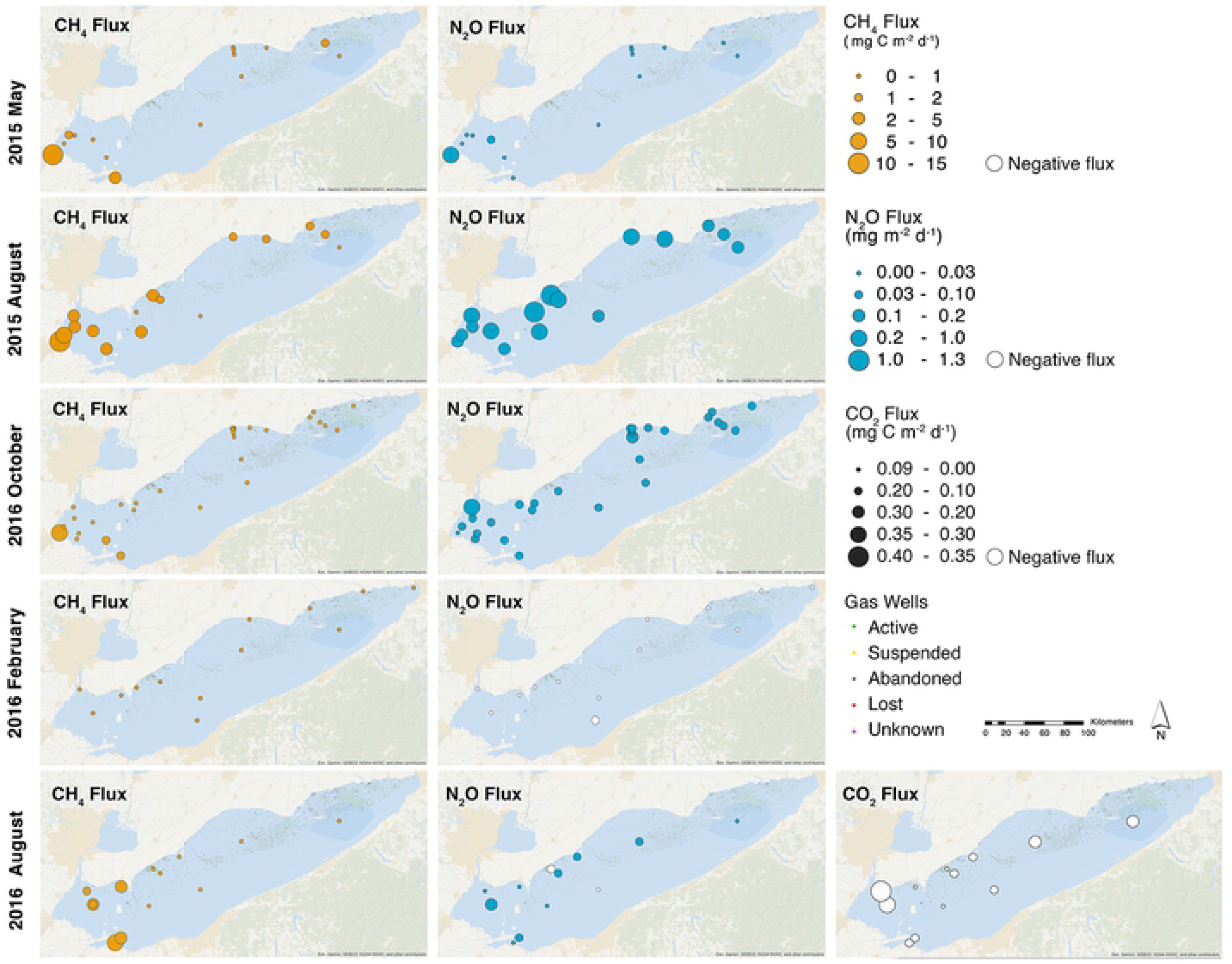
Sampling locations and methane (CH_4_), nitrous oxide (N_2_O), and carbon dioxide (CO_2_) fluxes in Lake Erie observed over the entire study period. CH_4_ fluxes are shown in the left column and the legend is the same for each month of the study period (mg C m^−2^ d^−1^). N_2_O fluxes are shown in the center column and both positive and negative fluxes are shown (mg N m^−2^ d^−1^). Positive N_2_O fluxes are shown in colors and negative (i.e., N_2_O uptake) in white. CO_2_ fluxes were only measured in August 2016 and are shown in mg C m^−2^ d^−1^; all CO_2_ fluxes were negative at this time. Also shown are the locations of natural gas wells and their status (active, abandoned, suspended, lost, or unknown).

Surface water samples for greenhouse gas (CH_4_ and N_2_O) analysis were taken at approximately at 1 m depth. Niskin bottles (5 L or 20 L) were used to collect surface water at 1 m depth into 155 mL glass serum bottles. Samples were overfilled three times using a tube at the bottom of each bottle to minimize gas loss and atmospheric contamination. Saturated HgCl_2_ brine (100 μL) was pipetted into the sample for preservation. Samples were sealed with gray butyl rubber septa and an aluminum crimp to minimize exposure to the atmosphere. Greenhouse gas samples were stored at room temperature until analysis.

Samples were also taken for pH, alkalinity, and dissolved organic carbon (DIC) concentration and isotopic (δ^13^C-DIC) analysis in August 2016. Surface water samples for pH analysis were transferred from Niskin bottles into clean 60 mL HDPE bottles as soon as the CTD rosette was brought onboard. These samples were poured into a clean 50 mL beaker with a magnetic stir rod and placed above a stir plate and pH was measured onboard using a Thermoscientific Orion 4-Star pH-Conductivity bench top meter with a refillable Ag/AgCl electrode. Water for alkalinity analysis was collected directly from the Niskin spout into two 60 mL syringes with an attachable 0.45 μm syringe filter. Alkalinity samples were filtered into a clean 125 mL HDPE bottle and either analyzed on ship after room temperature was reached or samples were stored at 4°C for later analysis.

Water samples for δ^13^C-DIC analysis were collected directly from the Niskin spout into a 30 mL syringe. The water was then filtered with an attachable Puradisc 0.7μm glass microfiber filter and delivered into a 2 mL wide mouth glass autosampler vial through a short 18 g needle. After the sample overflowed into the vial, it was then capped with an 11 mm PTFE rubber septa and crimped shut. The samples were then stored at 4°C.

### Sample Analysis

Dissolved gases were extracted from each sample using headspace extraction methods. At room temperature, 30 mL of ultra-high purity N_2_ was injected into a water sample in a serum vial, and 30 mL of water displaced into a second syringe, which created an inert headspace within the sample bottle. The bottles were then agitated for one minute with a Fisher Scientific Multi-Tube Vortexer to release any dissolved gases. As the 30 mL of displaced water was placed back into the sample bottle, a 30 mL syringe was used to extract the released gases from the headspace. The headspace gas was then transferred into a previously evacuated glass vial containing desiccant beads that was sealed with a butyl rubber septum and aluminum crimp cap. These headspace samples were analyzed using a Shimadzu Scientific Instruments Gas Chromatograph (GC)-2014 Greenhouse Gas Analyzer equipped with a flame ionization detector (FID) and an electron capture detector (ECD) at the University of Cincinnati.

Calibrated greenhouse gas standards were placed intermittently throughout the sample run to generate a calibration curve. Concentrations of calibrated CH_4_ standards range from 2.18 to 1,000 ppm in air (*n* = 4) and N_2_O standards range from 293 to 393 ppb (*n* = 3). The original dissolved gas concentrations in the water samples were determined with the temperature-specific Bunsen solubility coefficients (18) using the measured headspace gas concentrations. The equilibrium dissolved gas concentrations were calculated using water temperature, barometric pressure, and monthly average background atmospheric CH_4_ and N_2_O concentrations at the Mauna Loa Observatory during the time of sampling (28,29). Variation in final dissolved greenhouse gas concentration is about 6% using headspace extraction in our laboratory (13,14).

Alkalinity was measured by acid titration using a HACH Digital titrator equipped with sulfuric acid (0.16 N) and using a Thermoscientific Orion 4-Star pH-Conductivity bench top with a refillable Ag/AgCl electrode to record the inflection points. The recorded pH, acid count, and temperatures were entered into the USGS Alkalinity Calculator version 2.22 (30), which analyzes the titration curve and calculates the alkalinity using an acid correction factor of 1.01 and the inflection point method. DIC concentration and δ^13^C-DIC were measured via isotope ratio mass spectrometry at Skidaway Institute of Oceanography via the method of (31).

### Surface Flux Analysis

Surface fluxes of CH_4_ and N_2_O were calculated for all of the sampling periods using the following equation:

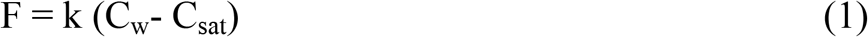

Where F is the flux, k is the gas transfer velocity, C_w_ is the measured CH_4_ or N_2_O concentration in surface water (measured as described above), and C_sat_ is the calculated equilibrium concentration of atmospheric CH_4_ or N_2_O in surface water (calculated as described above) (32). The gas transfer velocity (k) is calculated with the average wind speed and corresponding temperature dependent freshwater Schmidt number (33). Wind speed was averaged from the Port Stanley buoy, station 45132, for the duration of each sampling period (https://www.ndbc.noaa.gov/station_page.php?station=45132) (usually one week).

CO_2_ surface fluxes were calculated via USGS CO_2_-calc v4.0.9 (34). Measured values of pH, alkalinity, water temperature and average wind speed during sampling periods were used as the primary variables. Flux velocity is determined as a function of the average wind speed, and a Schmidt number of 600 was used for freshwater at 20°C (33). The CO_2_ constants (K_1_, K_2_, and K_w_) (35) and pressure effects (36) were chosen accordingly for freshwater. CO_2_ fluxes for each site (in August 2016 only) are shown in Table S6.

### Scaling Up

We categorized each site visited during each sampling trip to the Western, Central, or Eastern basin of Lake Erie based on its location within the lake and lake bathymetry (23) (Tables S1-S5). We then calculated monthly average CH_4_ and N_2_O emission rates for each basin for February, May, August (for each year and using an average of both 2015 and 2016 data), and October (Table S7 and S8). We then multiplied the average emission rate for each basin and each month by the surface area of each basin (Western basin = 3,284 km^2^; Central basin = 16,138 km^2^; Eastern basin = 6,235 km^2^), multiplied by the number of days in each month, and added up the emissions from each basin to get a monthly total (Tables S9 and S10). We used the average 2015 and 2016 emission rates for August for final calculations. Finally, we used linear interpolation to scale between measurement months to estimate an annual total for both CH_4_ and N_2_O (Fig S1).

### Error Estimation

To estimate the range in monthly and annual CH_4_ and N_2_O emissions, we calculated an average percent error for each gas. For each basin of the lake and each sampling month, we calculated the standard error of the average emission rate of CH_4_ or N_2_O (the standard deviation divided by the square root of the number of measurements). We then converted this to a percent error by dividing the standard error by the emission rate for that month. We calculated an average percent error for each basin and each gas, and then averaged each of those to estimate an average error for the emission measurement for CH_4_ and N_2_O, respectively. This method assumes that the largest source of variability in emission rates is differences within basins, and not measurement uncertainties – which are small compared to the spatial and temporal variability we observed in fluxes. The average percent error for CH_4_ was 19% and the average percent error for N_2_O was 37%. For some months, such as May 2015 in the Western Basin, we observed simultaneous high N_2_O fluxes in the Maumee River mouth and negative N_2_O fluxes elsewhere in the Western Basin, which leads to higher error estimates for N_2_O as compared to CH_4_. We also observed both negative and positive N_2_O fluxes in the Central Basin in May 2015 and August 2016.

## Results and discussion

### Methane

We found positive emissions of CH_4_ throughout the lake all year, including February (Fig 1). The highest CH_4_ emissions were in spring and summer, and from the shallow (Z_max_ ~10 m) (37) Western Basin (Figs 1 and 2). Emissions were lower in the deeper Central Basin, despite summer stratification, anoxia, and previous reports of CH_4_ buildup in the hypolimnion (18). The highest CH_4_ emissions from Lake Erie are similar to those from thawing permafrost lakes and ponds in the Arctic (38); however, unlike in Arctic lakes, we found low under-ice CH_4_ concentrations and emissions in winter (Figs 1 and 2).

**Fig. 2.**
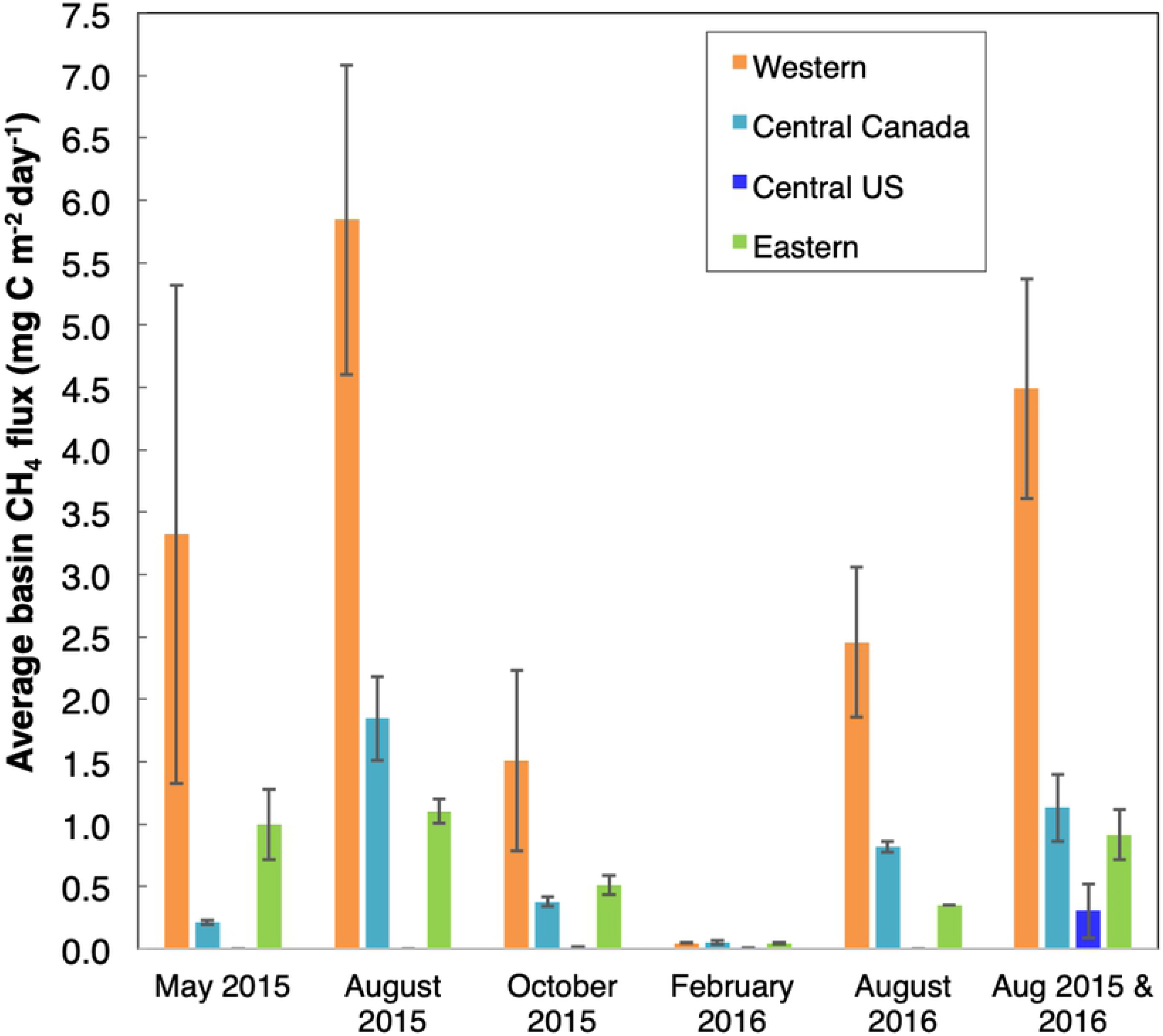
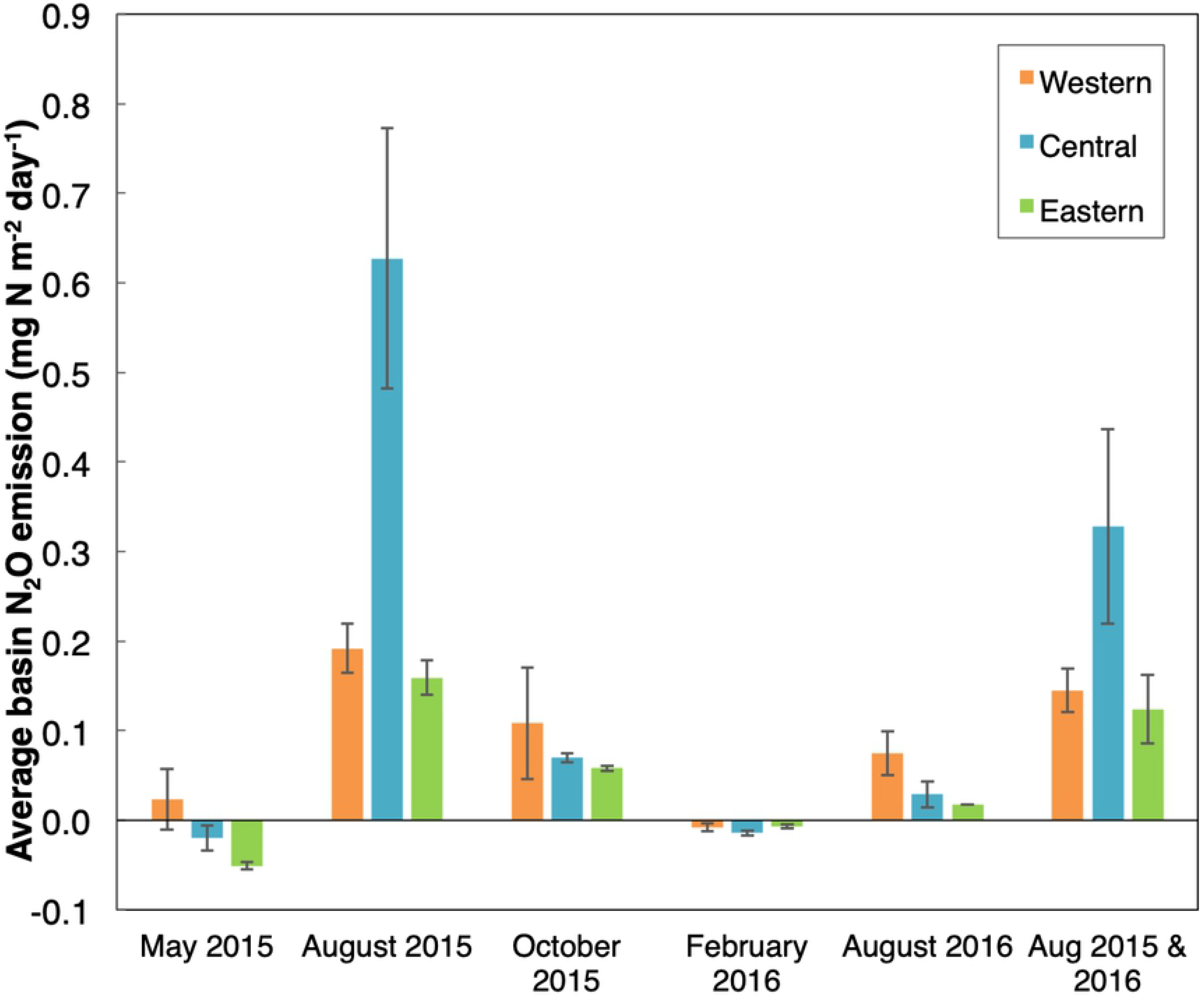
Basin-wide average CH_4_ (left) and N_2_O (right) fluxes for each month of measurements in the study. Also shown are separate averages for August 2015 and 2016 and both August trips together (Aug 2015 & 2016). Error bars represent standard error. CH_4_ plot shows separate columns for the Central Basin of Canada and US, as natural gas drilling is permitted in Canadian waters only.

Methane was previously measured in the Western Basin of Lake Erie in late summer 1969, near the Lake Erie islands (17)(39). The 1969 emission rate was 0.5 mg C m^−2^ d^−1^, ~10 times lower than our measurement from approximately the same location in August 2015 (site 478, 4.6 mg C m^−2^ d^−1^; Table S2) (17)(39). These data suggest that CH_4_ emissions from Western Lake Erie have increased. The (17) reference concluded that “methane…must be considered in the total cycle of carbon in this eutrophic lake” (17). Despite this important observation, however, no further measures of CH_4_ were published until our recent study in 2016.

Contrary to our previous work using stable isotopes (18), the current data do not indicate that natural gas production is a major source of CH_4_ in Lake Erie, for several reasons. In the current study, we measured CH_4_ emissions in winter, when temperature-dependent biological CH_4_ production is expected to be lowest, and found no significant difference between CH_4_ emissions in the Central and Eastern basin over gas wells (average flux = 0.05 ± 0.004 mg C m^−2^ d^−1^) versus those not (0.04 ± 0.003 mg C m^−2^ d^−1^). We also found no difference in δ^13^C-DIC or DIC concentration in samples taken in bottom waters in August 2016 in over gas wells versus non-gas producing areas (see further discussion below). Furthermore, the current study showed that the majority of CH_4_ emissions were largely found in the Western Basin, not sampled in our previous study, where there are no gas wells (Fig 1). Emissions of CH_4_ from the Western Basin are most likely derived from biogenic CH_4_ production in sediments. Finally, we note that most wells in Lake Erie are plugged and not producing natural gas, and thus would not be expected to be a large CH_4_ source (40).

A recent study reported CH_4_ concentrations and isotopic composition of CH_4_ in Lake Michigan and Lake Superior from June 2017 (21). Methane concentrations in summer in Lake Erie were more than 200 times higher than in Lake Superior (21) (Table S1). Surface water CH_4_ concentrations ranged from 3.5 to 60 nM in Lakes Michigan and Superior (21), much lower than in Lake Erie, where concentrations in May 2015 (the closest sampling month to June) ranged from 14 to 780 nM (Table S1). The highest CH_4_ concentrations in (21) were found in coastal Lake Michigan where the highest primary production rates are found, and the highest sediment methanogenesis rates are likely (21), similar to the current study. As we also concluded in the current study, radiocarbon measurements indicated no input of CH_4_ from old sources, such as fossil fuels (21), although radiocarbon measurements of CH_4_ in Lake Erie would help further indicate the relative inputs of natural gas and biogenic CH_4_ in active production areas.

### Nitrous Oxide

N_2_O emissions showed different spatial and temporal patterns than CH_4_ (Figs. 1 and 2). This is not surprising, as CH_4_ is likely produced in anoxic sediments whereas N_2_O production and consumption occur under both aerobic and anaerobic conditions in sediments and the water column (41). The highest N_2_O emissions were in the Central Basin in August 2015 (Fig 1). N_2_O emissions also differed between the two August sampling trips (Figs 1 and 2): in the north-central basin, higher emissions were observed in 2015 than in August 2016, when a phytoplankton bloom likely reduced dissolved N concentrations and therefore nitrification. We also observed some negative N_2_O fluxes in the Central and Western Basin in May 2015 and August 2016, and throughout the lake in February 2016 (Figs 1 and 2).

There are no recent studies of N_2_O in Lake Erie. In August 1977, a negative N_2_O flux was measured near the mouth of the Detroit River (19), in comparison to our fluxes of 0.4 mg N m^−2^ d^−1^ in August 2015 (site 881; Table S2) and 0.02 mg N m^−2^ d^−1^ in August 2016 (site 971; Table S5) in the same area (Fig. 1). N_2_O emissions have increased in Lake Erie: indeed, our data indicate that western Lake Erie has flipped from an overall sink to a source of N_2_O along with re-eutrophication. Eutrophic lakes are not currently considered a source of anthropogenic N_2_O in global budgets (42), although recent studies suggest their emission rate is globally significant (10).

Nitrous oxide uptake has been documented previously in eutrophic reservoirs and agricultural lakes. A recent study of eutrophic agricultural reservoirs in the Great Plains of Canada found widespread undersaturation and uptake of N_2_O (43). A study of a eutrophic reservoir near Cincinnati, Ohio found that denitrification consumed N_2_O at some periods during the year although it was an N_2_O source on annual time scales (13). Overall, there are less studies that have included N_2_O than CH_4_ in eutrophic lakes and reservoirs (43), and these studies indicate that more measurements are needed to elucidate the role of N_2_O in an increasingly warmer, wetter, and nutrient enriched world.

### Carbon Dioxide

Carbon dioxide emission rates were only measured in August 2016, and all fluxes were negative, reflecting high photosynthesis and/or low respiration (Fig 1). Methane oxidation contributes to the dissolved inorganic C pool. Methane is ^13^C-depleted relative to CO_2_ (44); we observed lower δ^13^C-DIC concentrations at sites with higher DIC and CH_4_ concentrations in bottom waters throughout Lake Erie (Fig. 3, Table S6), indicating a greater presence of DIC from CH_4_ oxidation in samples with high DIC (Fig 3). There was no difference in δ^13^C-DIC in sites with and without gas wells, indicating biogenic CH_4_ is the predominant source of CH_4_ in Lake Erie (Fig 3), in contrast to our previous study (18). These data indicate that if positive CO_2_ emissions occur later in the season, the production of CH_4_ in the lake due to eutrophication may enhance biogenic CO_2_ emissions. CO_2_ is a large contributor to the GHG footprint of other large eutrophic lakes (10). However, it is unknown whether Lake Erie is a net sink or source for CO_2_ on annual time scales (45)(46), and thus whether it contributes to increasing atmospheric CO_2_ and climate warming. The current and predicted CO_2_ balance of large lakes such as Lake Erie is a critical data gap that should be targeted by future work (47).

**Fig 3.**
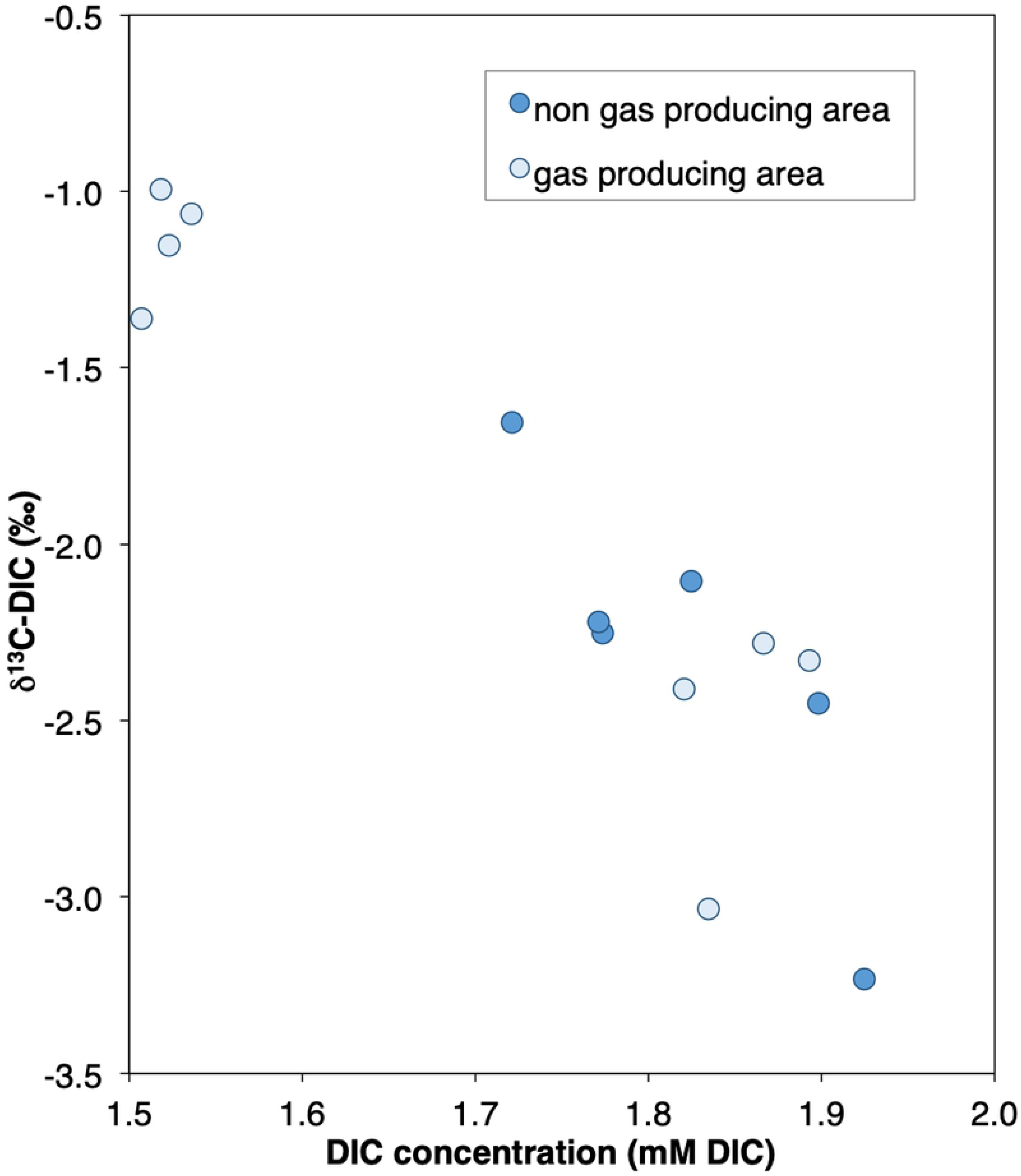
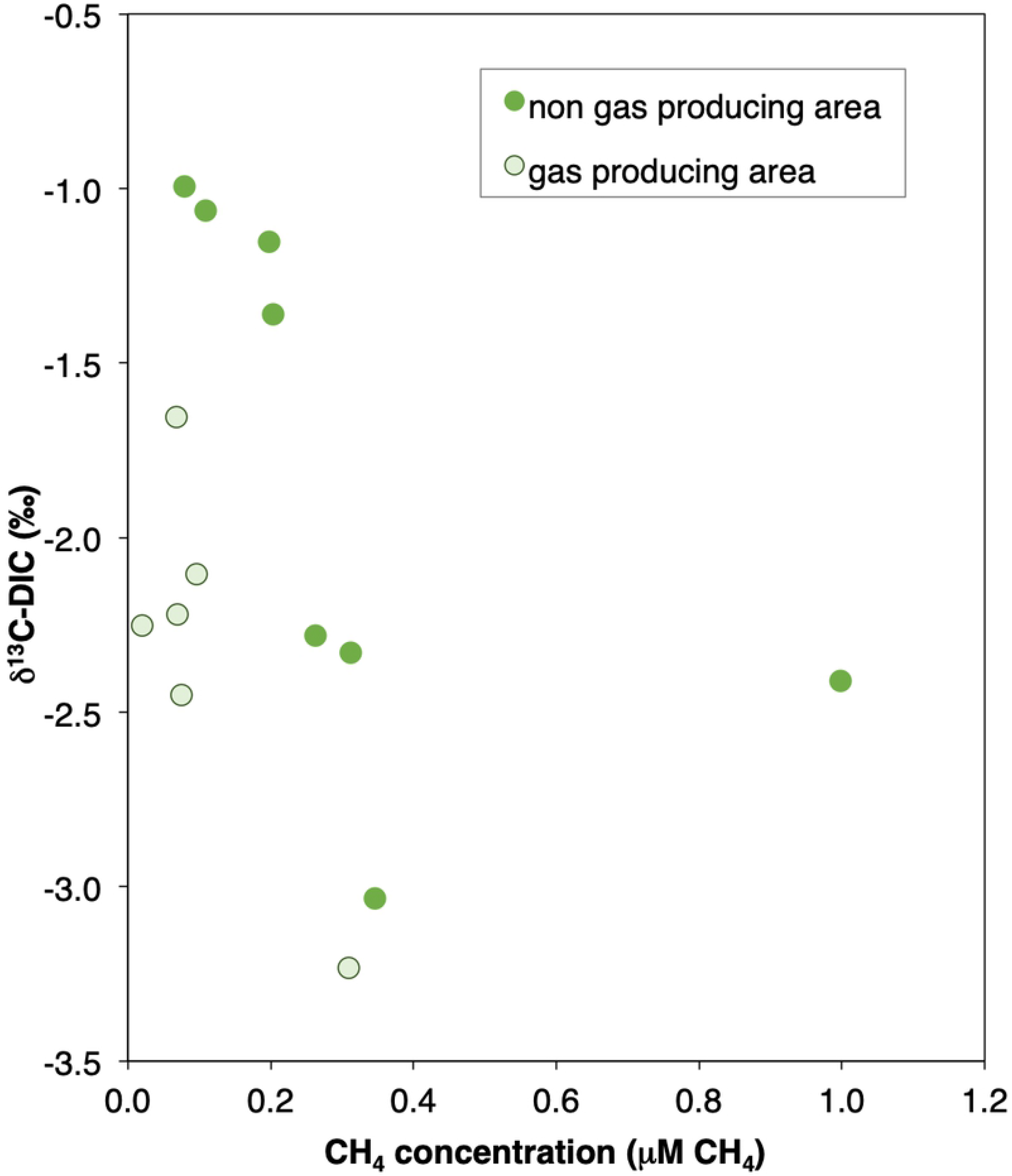
Carbon isotopic composition (δ^13^C [‰; VPDB]) of dissolved inorganic carbon (DIC) versus the DIC concentration (left) and the CH_4_ concentration (right). All data are from samples taken 1 m from the bottom sediments. Samples taken proximal to and away from natural gas wells are indicated.

### Scaling Up

CH_4_ and N_2_O emissions occur most in spring, summer, and fall in Lake Erie (Fig. 2), and most in the Western Basin (particularly for CH_4_), corresponding with highest seasonal phytoplankton biomass (May-October; (2)). We estimate annual emissions from Lake Erie of 6,294 ± 1,259 metric tons CH_4_-C yr^−1^ and 597 ± 209 metric tons N_2_O-N yr^−1^, or 209,775 metric tons CO_2_-equivalent (CO_2_-e) for CH_4_ and 279,286 metric tons CO_2_-e for N_2_O (16)(48). This is the equivalent of over 100,000 cars driven for one year, or almost 250 million kilograms of coal burned in a year (48).

We compared our results to data published by USEPA on CH_4_ emissions in the Great Lakes region from large facilities in the waste, oil and natural gas, and fertilizer production sectors (49). This showed that Lake Erie emits CH_4_ at a rate comparable to and higher than most landfills and natural gas distribution systems in the US states of Ohio and Michigan. By way of comparison, in 2017 over 200 facilities reported emissions in Ohio, and only three landfills, one natural gas distribution company, and one fertilizer plant had CH_4_ emissions greater than our annual estimate for Lake Erie, while in Michigan fewer than six of the 200+ facilities reporting CH_4_ exceeded those of Lake Erie. Furthermore, Lake Erie represented a larger N_2_O source than any other industrial source in Ohio or Michigan reported to EPA in 2017, although, fertilized agricultural soils, the largest anthropogenic N_2_O source regionally and nationally (42), do not appear in this database.

## Conclusions

We present a new and significant observation on the negative effects of phytoplankton blooms in Lake Erie. If climate change and eutrophication increase CH_4_ and N_2_O emissions from Lake Erie, as indicated here and elsewhere (10)(22)(11), this represents a positive feedback to climate warming. A warmer and wetter climate may lead to higher CH_4_ and N_2_O emissions from Lake Erie, particularly with increasing nutrient inputs. Carbon cycle-climate change feedbacks are the main driver of interannual variability and increases in global CH_4_ concentrations (42)(50). A growing body of evidence also shows that eutrophic water bodies such as Lake Erie also contribute to the global N_2_O budget with a positive feedback to climate warming (42). These results also have significant implications for other inland waters and imply that greenhouse gas emissions from eutrophic lakes – particularly large ones – should be considered for inclusion in anthropogenic greenhouse gas budgets, as these represent regionally significant fluxes.

## Acknowledgments

The authors thank F. Åkerström, C. Botner, D. Disbennett, A. Fries, and K. Jimenez for laboratory assistance; J. Beaulieu, R. Bourbonierre, M. Griffin, and C. Winslow for helpful discussions while designing this study; and the captain and crew of the *CCGS Limnos* and the *CCGS Griffon* for sampling assistance and for keeping us alive and supplied with poutine during research cruises.

## Author contributions

J.M.F. and A.T.S. designed the study and interpreted results with input from all authors; J.M.F. collected the samples and analyzed them for CH_4_, N_2_O, and CO_2_; J.A.B. analyzed samples for DIC and DIC isotopes; and J.M.F. and A.T.S. wrote the paper with contributions from all authors.

## Data and materials availability

All data and methods needed to reproduce the results in the paper are provided in the supporting information.

## Supporting Information

**Table S1. May 2015 Lake Erie dissolved [CH_4_] and [N_2_O] and surface fluxes.** CCGS Limnos cruise dates were May 12th - 14th 2015. Data from water collected 1 meter below the surface water and atmosphere interface. Average windspeed was 4.73 m s^−1^. Sample locations are shown in Figure 1.

**Table S2. August 2015 Lake Erie dissolved [CH_4_] and [N_2_O] and surface fluxes.** CCGS Limnos cruise dates were August 24th - 27th 2015. Data from water collected 1 meter below the surface water and atmosphere interface. Average windspeed was 6.65 m s^−1^. Sample locations are shown in Figure 1.

**Table S3. October 2015 Lake Erie dissolved [CH_4_] and [N_2_O] and surface fluxes.** CCGS Limnos cruise dates were October 5th - 8th 2015. Data from water collected 1 meter below the surface water and atmosphere interface. Average windspeed was 3.84 m s^−1^. Sample locations are shown in Figure 1.

**Table S4. February 2016 Lake Erie dissolved [CH_4_] and [N_2_O] and surface fluxes.** CCGS Griffon cruise dates were February 15th - 19th 2016. Data from water collected 1 meter below the surface water and atmosphere interface. Average windspeed was 1.97 m s^−1^. Sample locations are shown in Figure 1.

**Table S5. August 2016 Lake Erie dissolved [CH_4_] and [N_2_O] and surface fluxes.** CCGS Limnos cruise dates were August 29th to September 1^st^ 2016. Data from water collected 1 meter below the surface water and atmosphere interface. Average windspeed was 4.05 m s^−1^. Sample locations are shown in Figure 1.

**Table S6. August 2016 Lake Erie [DIC], ^13^C-DIC, [CO_2_], CO_2_ surface flux and relevant sample information.** CCGS Limnos cruise dates were August 29th to September 1^st^ 2016. Data from water collected 1 meter below the surface water and atmosphere interface, and from 1 m from the sediment water interface. Average windspeed was 4.05 m s^−1^. DIC concentration and ^13^C-DIC was determined via isotope ratio mass spectrometry at Skidaway Institute of Oceanography via the method of Brandes (2009). Sample locations are shown in Figure 1.

**Table S7. Monthly average CH_4_ surface fluxes for each basin of Lake Erie.**

**Table S8. Monthly average N_2_O surface fluxes for each basin of Lake Erie.**

**Table S9. Weighted average CH_4_ flux of each sampling month of the whole lake.** The daily average flux is multiplied by the surface area (Bolsenga and Herdenforf, 1993) of the designated basin for an estimate of the total flux of the basin. The sum of all the basins for each sampling period is the estimated daily flux of the whole lake. Multiplying the lake total by the number of days in the sampling month estimates the total flux for the month.

**Table S10. The weighted average N_2_O flux of each sampling month of the whole lake.** The daily average flux is multiplied by the surface area (Bolsenga and Herdenforf, 1993) of the designated basin for an estimate of the total flux of the basin. The sum of all the basins for each sampling period is the estimated daily flux of the whole lake. Multiplying the lake total by the number of days in the sampling month estimates the total flux for the month.

**Table S11. Average annual CH_4_ and N_2_O surface flux.** A monthly average was produced for the months that were not measured by calculating the linear regression between each measured month and then applying this regression to the months between. The annual total for both CH_4_ and N_2_O is the sum of the calculated and estimated values for a full year. These measurements and estimates are also shown in Figure S1.

**Fig. S1. Annual trends in average lake-wide emissions of CH_4_ (left) and N_2_O (right).** The larger points on each curve represent months when measurements were made (Fig. 1), and smaller points are estimated using linear extrapolation as described in the methods. The lake is a positive source of CH_4_ all year, and a weak sink for N_2_O in winter, but fluxes of both gases are highest in late summer. Annual emissions were estimated by summing both measured and estimated monthly fluxes. For a discussion of error estimates, please see the supplemental methods.

